## Supplemental Figures and Tables for "Homodecameric Rad52 promotes single-position Rad51 nucleation in homologous recombination"

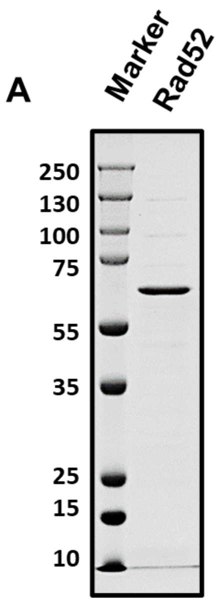

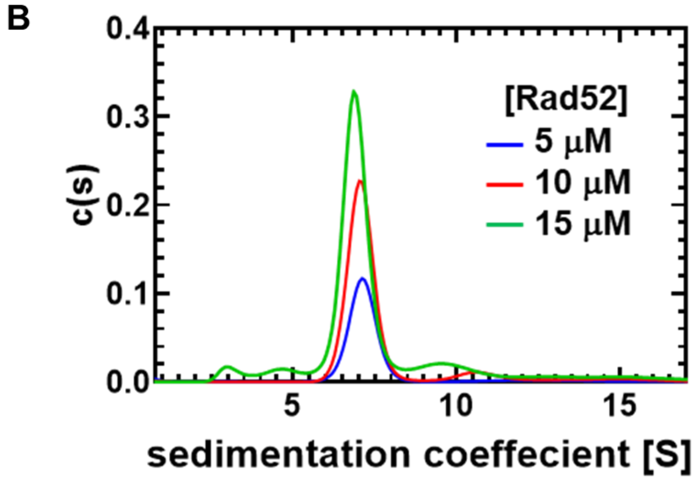

C

**AUC Sedimentation Velocity c(s) values**

| [Rad52] | sw | Sw (20,w) | Mw (kDa) | Stokes Radius 20°C (nm) | a/b (oblate) | a/b (prolate) | % |
|---|---|---|---|---|---|---|---|
| 5 μM | 7.25 | 11.3 | 532 | 11.2 | 19.0 | 16.7 | 73.7 |
| | 11.0 | 17.1 | 991 | 13.8 | 19.0 | 16.7 | 04.4 |
| | 14.4 | 22.4 | 1484 | 15.8 | 19.0 | 16.7 | 02.2 |
| 10 μM | 7.14 | 11.1 | 521 | 11.2 | 19.0 | 16.8 | 81.8 |
| | 10.6 | 16.5 | 944 | 13.6 | 19.0 | 16.8 | 05.9 |
| | 13.9 | 21.7 | 1421 | 15.6 | 19.0 | 16.8 | 04.2 |
| 15 μM | 7.46 | 11.6 | 537 | 11.0 | 17.9 | 15.7 | 70.8 |
| | 13.11 | 20.4 | 1250 | 14.6 | 17.9 | 15.7 | 3.0 |

**Supplementary Figure 1. *Saccharomyces cerevisiae* Rad52 is predominantly a single homodecameric species solution**. **A)** SDS-PAGE analysis of recombinantly overproduced *S. cerevisiae* Rad52. **B)** Sedimentation velocity analytical ultracentrifugation ($AUC^{SV}$) analysis of full-length *S. cerevisiae* Rad52 was performed at various concentrations of protein (monomer concentrations noted). Rad52 is predominantly a single species in solution with an estimated molecular mass corresponding to a homodecamer. **C)** The table below denotes the c(s) values calculated for each of the species and their relative abundance (%).

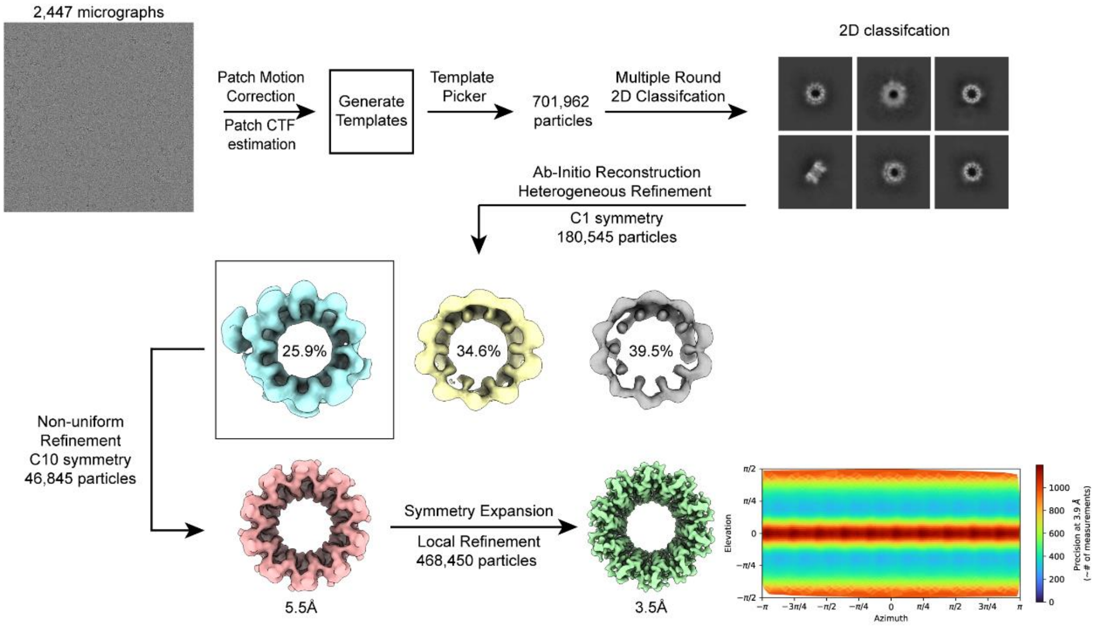

**Supplementary Figure 2. Workflow of single particle analysis of the yeast Rad52 Cryo-EM data.** Micrographs collected during imaging of Rad52 was analyzed as shown.

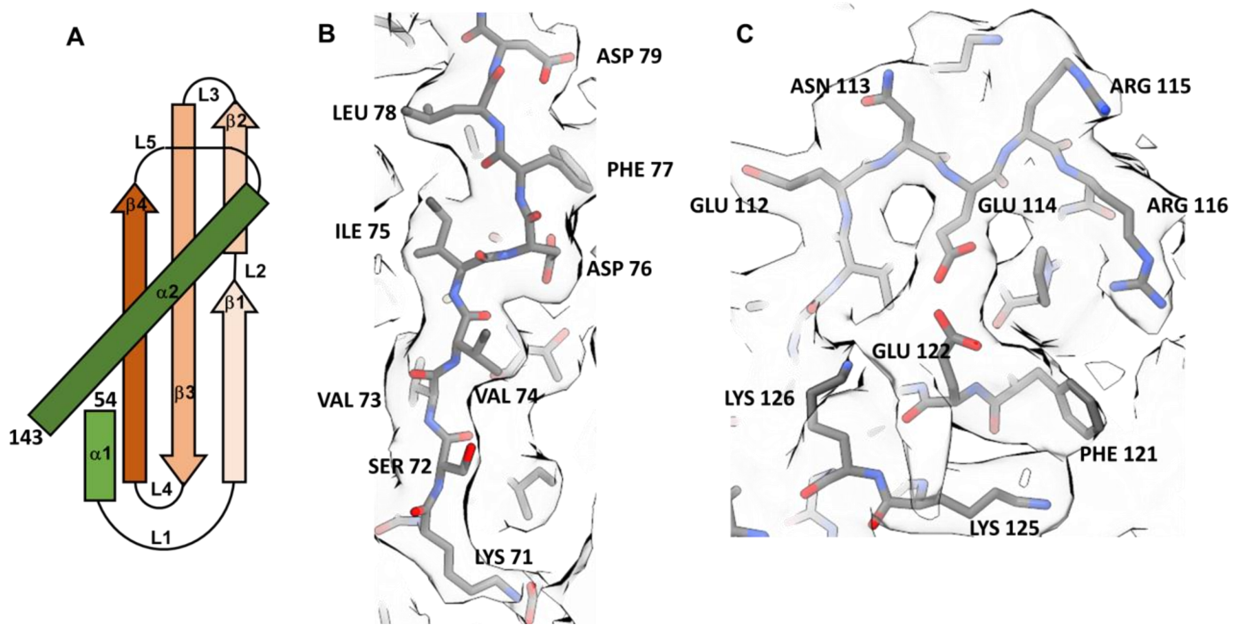

**Supplementary Figure 3. Structural details of *S. cerevisiae* Rad52. A)** Cartogram of the Rad52 monomer observed in the Cryo-EM structure. **B)** & **C)** Examples of sample maps and models of yeast Rad52.

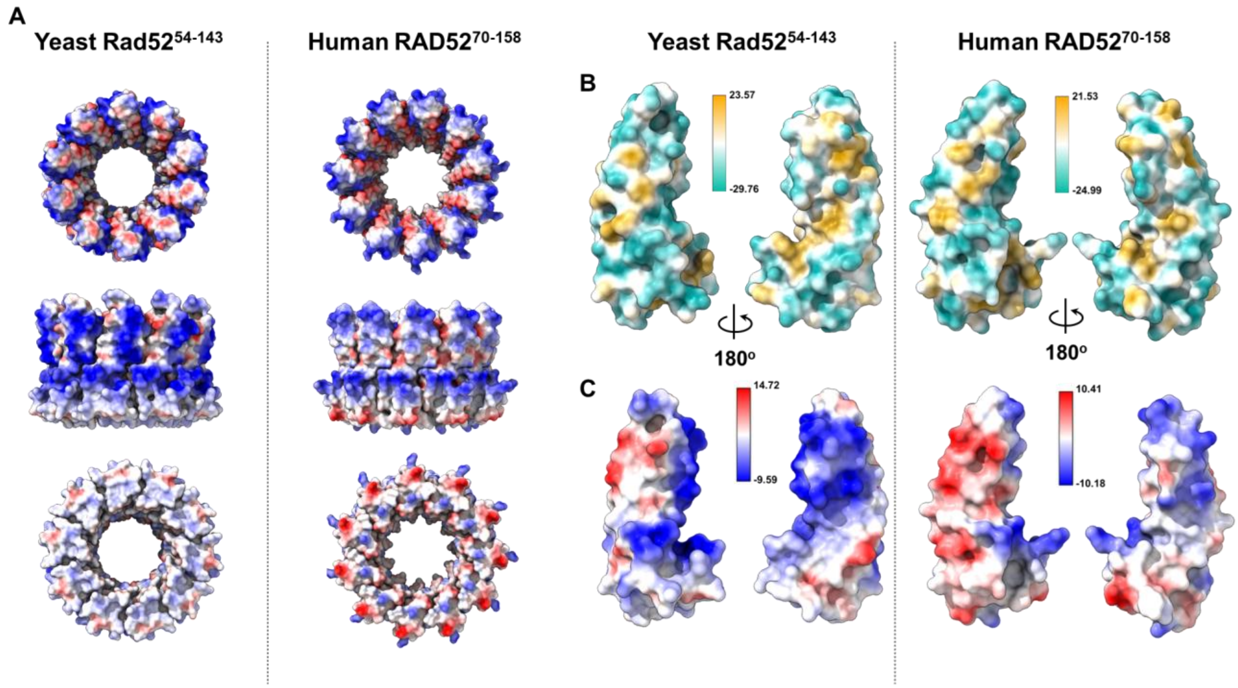

**Supplementary Figure 4. Comparison of yeast and human Rad52 structures. A)** A comparison of the electrostatic maps between yeast Rad52 and human RAD52 (PDB: 1KN0). Comparison of **B)** hydrophobicity and **C)** electrostatic maps within a single subunit between yeast Rad52 and the corresponding residues in the human RAD52. The comparisons reveal that there are additional positively charged regions in yeast Rad52 that are exposed and likely important for DNA interactions.

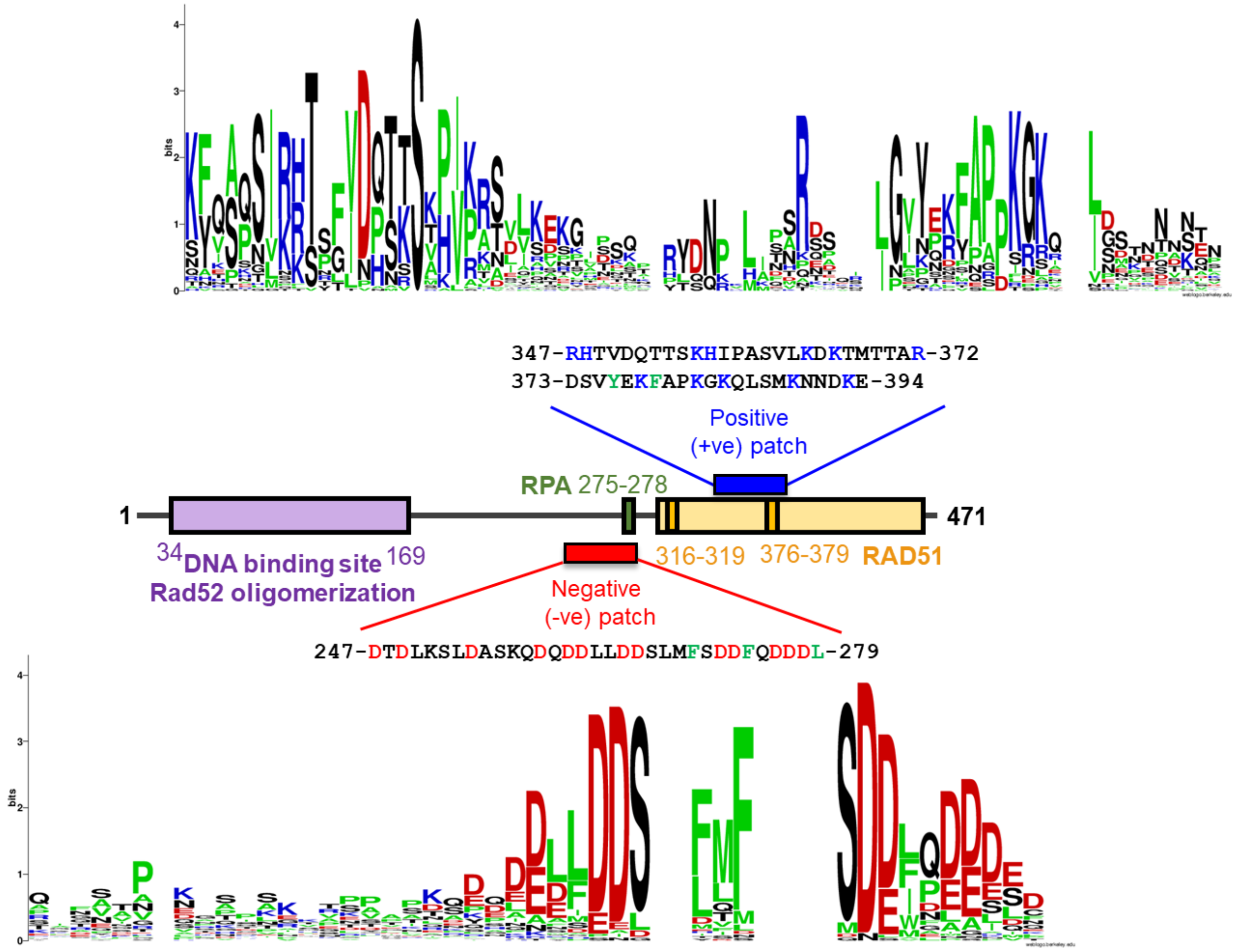

**Supplementary Figure 5. Sequence conservation analysis of the C-tail.** WebLogo represents sequence similarity analysis of regions in the disordered C-terminal half of Rad52 carrying the Rad51 binding site (positive patch) and the RPA binding site (negative patch). The analysis encompasses Rad52 proteins from 66 different fugal species. Several residues are highly conserved.

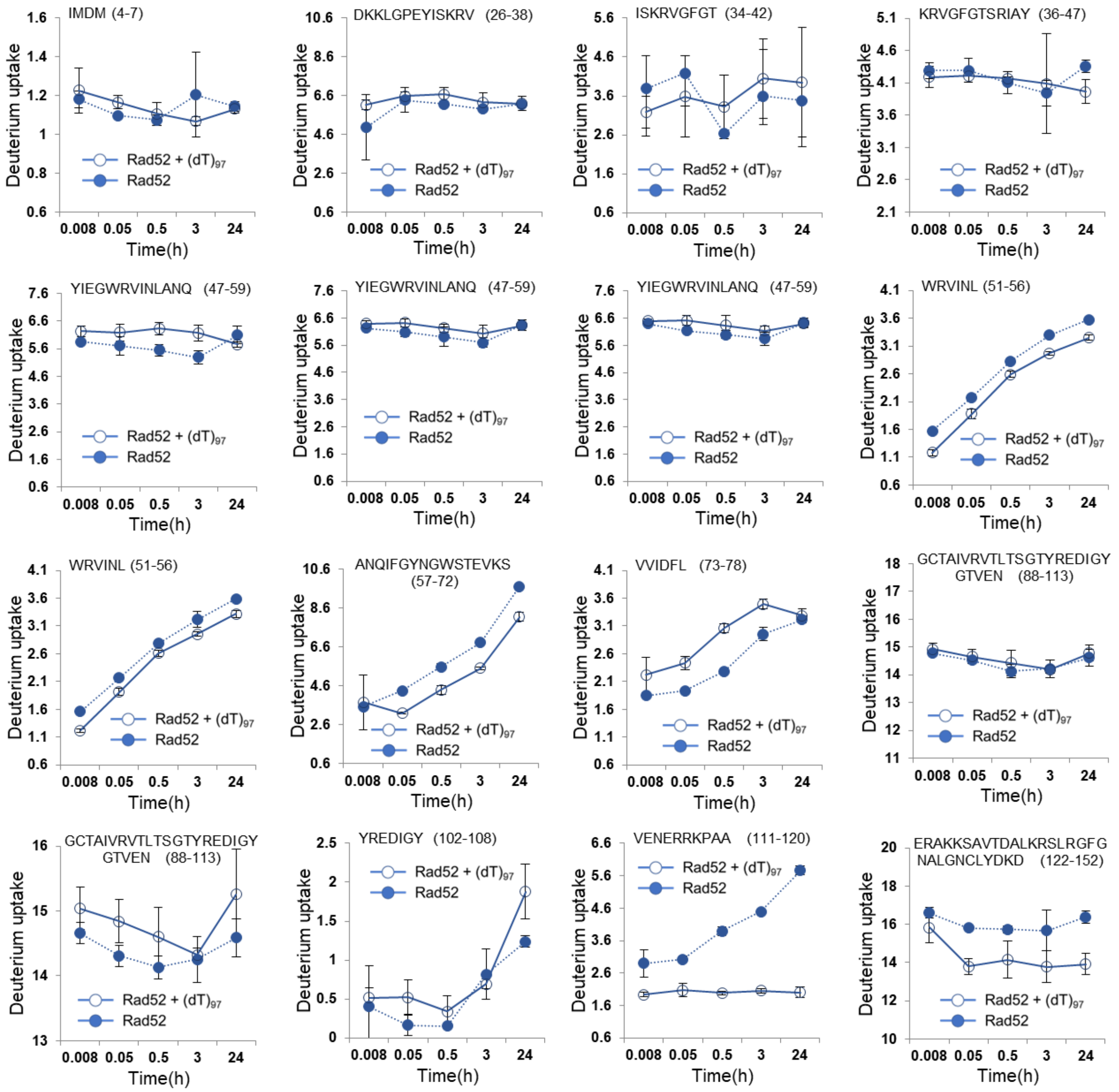

**Supplementary Figure 6. Deuterium uptake changes captured in HDX-MS analysis of yeast Rad52.** The plotted data for deuterium incorporation as a function of time are shown for individual peptides identified in the HDX-MS analysis. This panel depicts peptides corresponding to amino acid positions 1-152 in the N-terminal half. Data from both Rad52 and the Rad52-$(dT)_{97}$ complex are compared for each peptide.

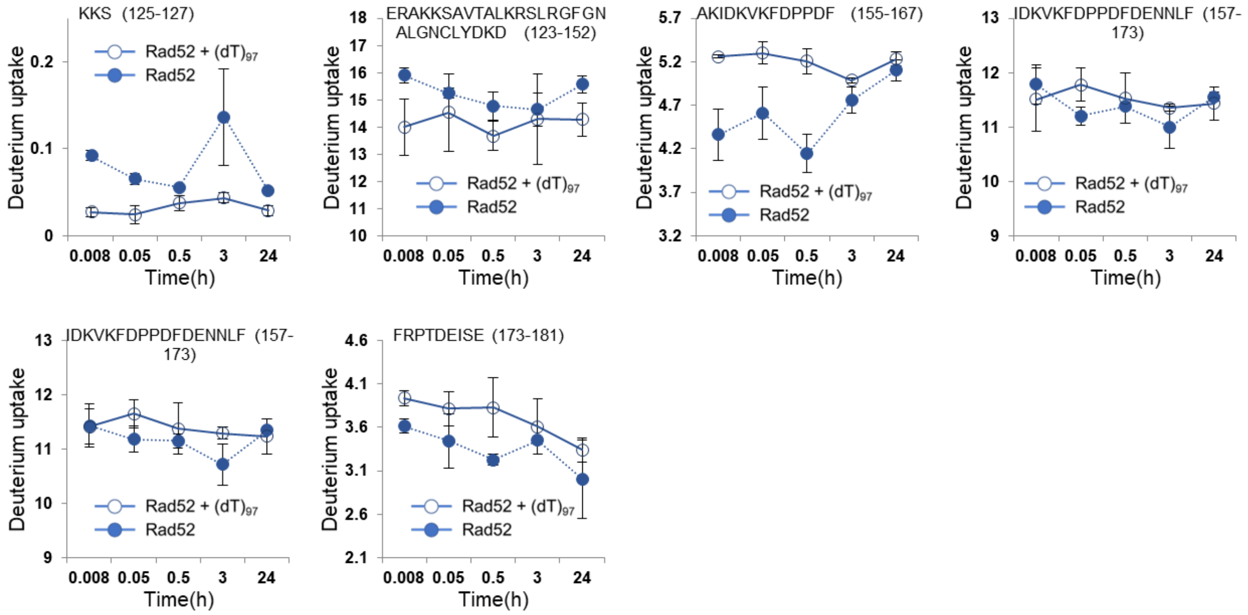

**Supplementary Figure 7. Deuterium uptake changes captured in HDX-MS analysis of yeast Rad52.** The plotted data for deuterium incorporation as a function of time are shown for individual peptides identified in the HDX-MS analysis. This panel depicts peptides corresponding to amino acid positions 125-181 in the N-terminal half. Data from both Rad52 and the Rad52-$(dT)_{97}$ complex are compared for each peptide.

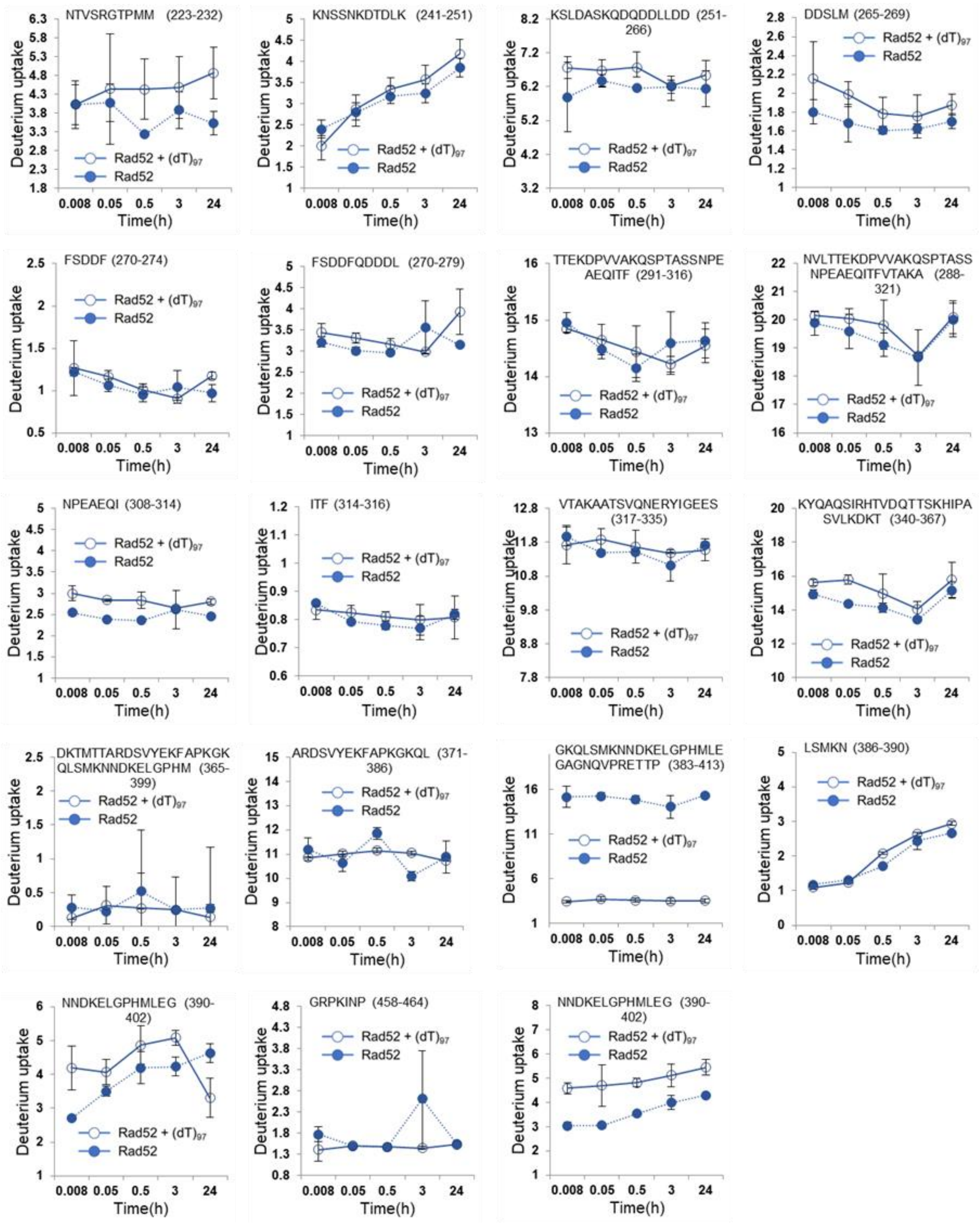

**Supplementary Figure 8. Deuterium uptake changes captured in HDX-MS analysis of yeast Rad52.** The plotted data for deuterium incorporation as a function of time are shown for individual peptides identified in the HDX-MS analysis. This panel depicts peptides corresponding to amino acid positions 223-464 in the C-terminal half. Data from both Rad52 and the Rad52-(dT)$_{97}$ complex are compared for each peptide.

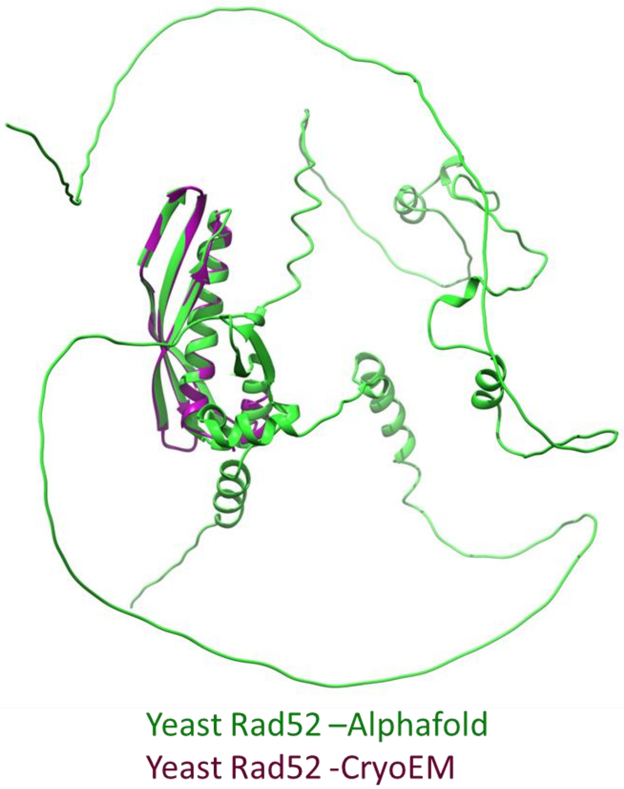

**Supplementary Figure 9. Comparison of the Alphafold model and cryo-EM structure of yeast Rad52.** The Alphafold model of *S. cerevisiae* Rad52 (AF-P06778-F1; green) overlaid with the ordered part of the N-terminal half observed in the cryo-EM structure (purple). Excellent agreement in the structures is observed in the N-terminal half between the two.

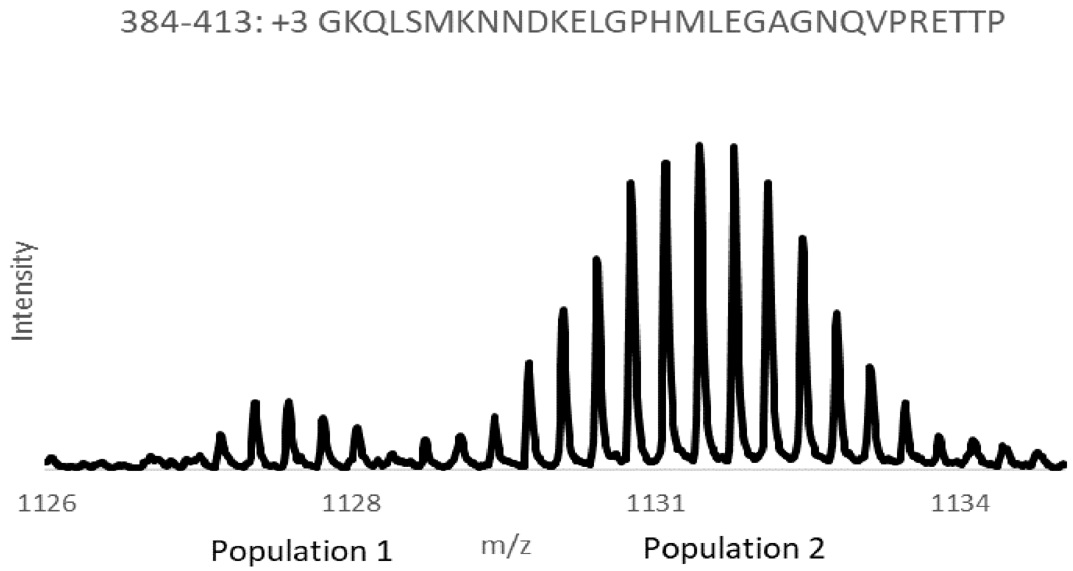

**Supplementary Figure 10. Hydrogen/deuterium exchange data of Rad52 peptide 384-413 has a bimodal distribution.** Mass spectra of the $(M+3H)^{3+}$ charge state of peptide 384-413 following incubation in 90% deuterium buffer for 30 seconds. Two distinct populations (envelopes) of deuterium uptake were observed. Relative abundance of the different peptide forms is shown in Fig 3E and 3F.

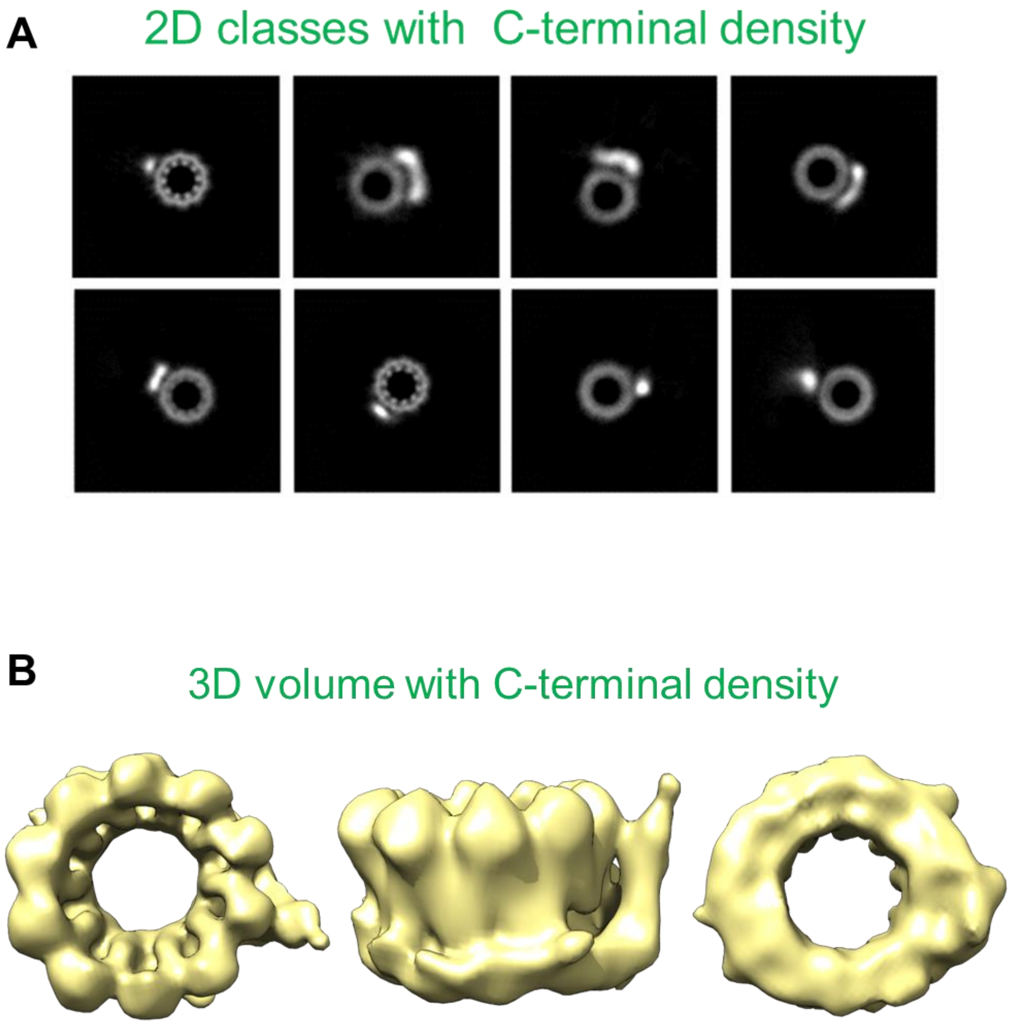

**Supplementary Figure 11. Additional density for C-terminal region positioned adjacent to the ordered N-terminal homodecameric ring. A)** Examples of 2D classes that are captured in the cryo-EM analysis of Rad52. These classes were discarded in the structure determination, but clearly show presence of additional density adjacent to the ordered N-terminal ring. **B)** Subsequent 3D volumes show that the C-terminal region sits adjacent to the ring but appears to be asymmetrically positioned to one part. The underlying allosteric mechanisms that prevent other subunits in the ring from interacting with their respective C-terminal regions remains to be determined.

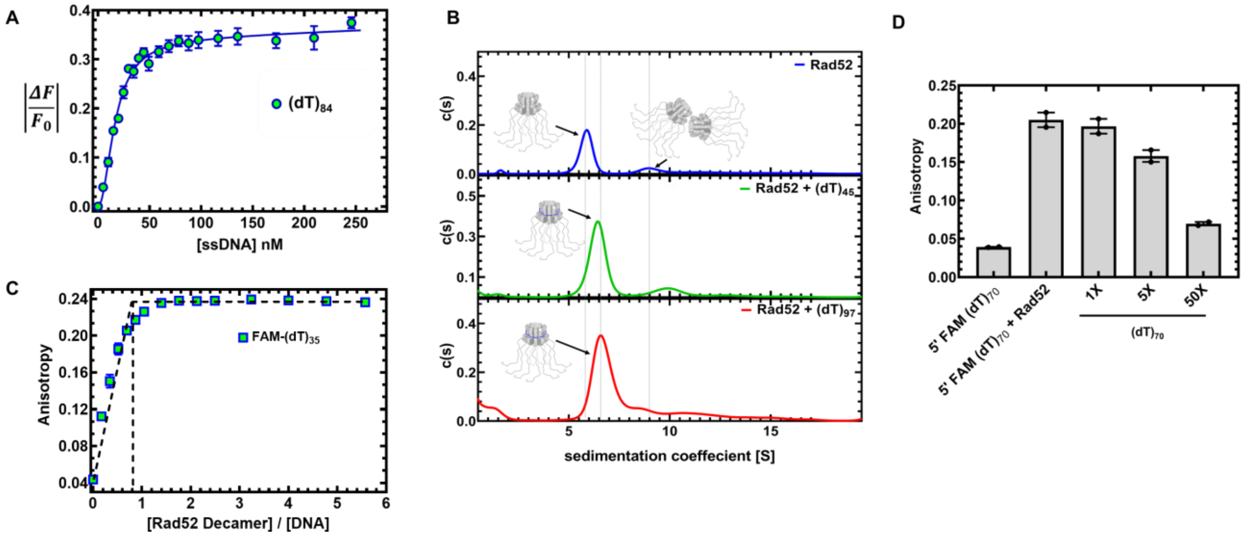

**Supplementary Figure 12. Rad52 binds to ssDNA as a homodecamer. A)** Intrinsic tryptophan quenching titration-based analysis of ssDNA $(dT)_{84}$ binding to Rad52 is consistent with two binding sites. The data are well described by a model describing two sequential binding sites: high affinity ($K_D$=1.3x10$^{-12}$M) and lower affinity binding site ($K_D$=28x10$^{-9}$M). **B)** Analytical ultracentrifugation analysis of Rad52 in the absence or presence of short [$(dT)_{45}$] or longer [$(dT)_{97}$] ssDNA substrates predominantly show a decamer forming 1:1 complex with DNA. The double decamer species is also capable of binding to ssDNA. **C)** Fluorescence anisotropy measurement of Rad52 binding to a 5′-FAM-$(dT)_{35}$ oligonucleotide shows formation of a stoichiometric complex. As expected, $(dT)_{35}$ is long enough to saturate only the high affinity site. **D)** Competition experiments were performed by titrating in increasing concentrations of unlabeled $(dT)_{70}$ into a preformed Rad52:5′-FAM-$(dT)_{70}$ complex. Data shows that Rad52 possesses a limited number of ssDNA binding sites, and the bound ssDNA is in dynamic equilibrium with the pool of free ssDNA in the solution.

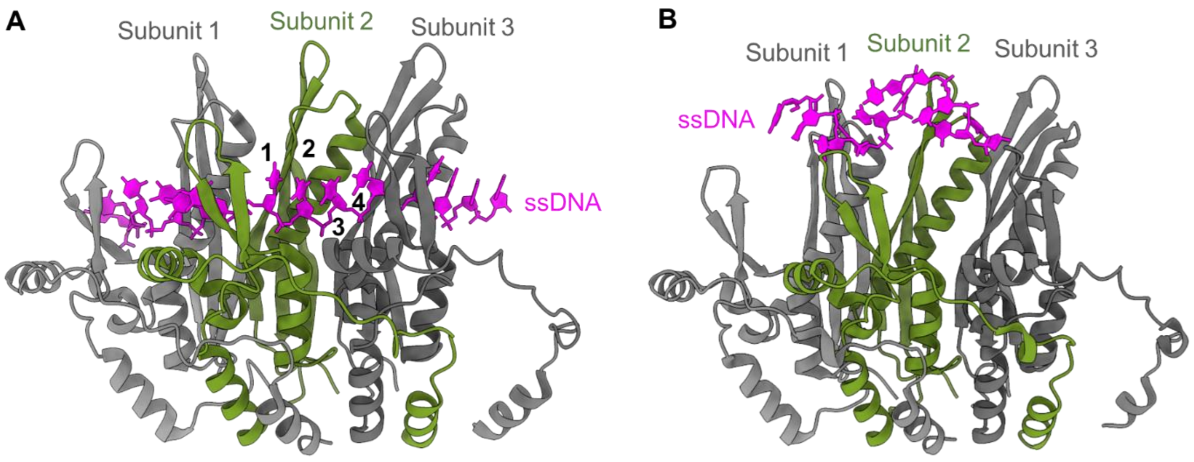

**Supplementary Figure 13. Crystal structures of ssDNA bound human RAD52.** Three subunits (colored individually) from the undecameric human RAD52 crystal structures are shown. **A)** ssDNA $(dT)_{40}$ bound to the inner binding site was captured when the outer binding site was perturbed by mutating Lys-102 and Lys-133 (PDB ID 5XRZ). **B)** 6 nt of a $(dT)_{40}$ ssDNA substrate is observed bound to outer site was captured sandwiched between two RAD52 undecameric rings (PDB ID 5XS0). Shown here are contacts between a single RAD52 ring and ssDNA.

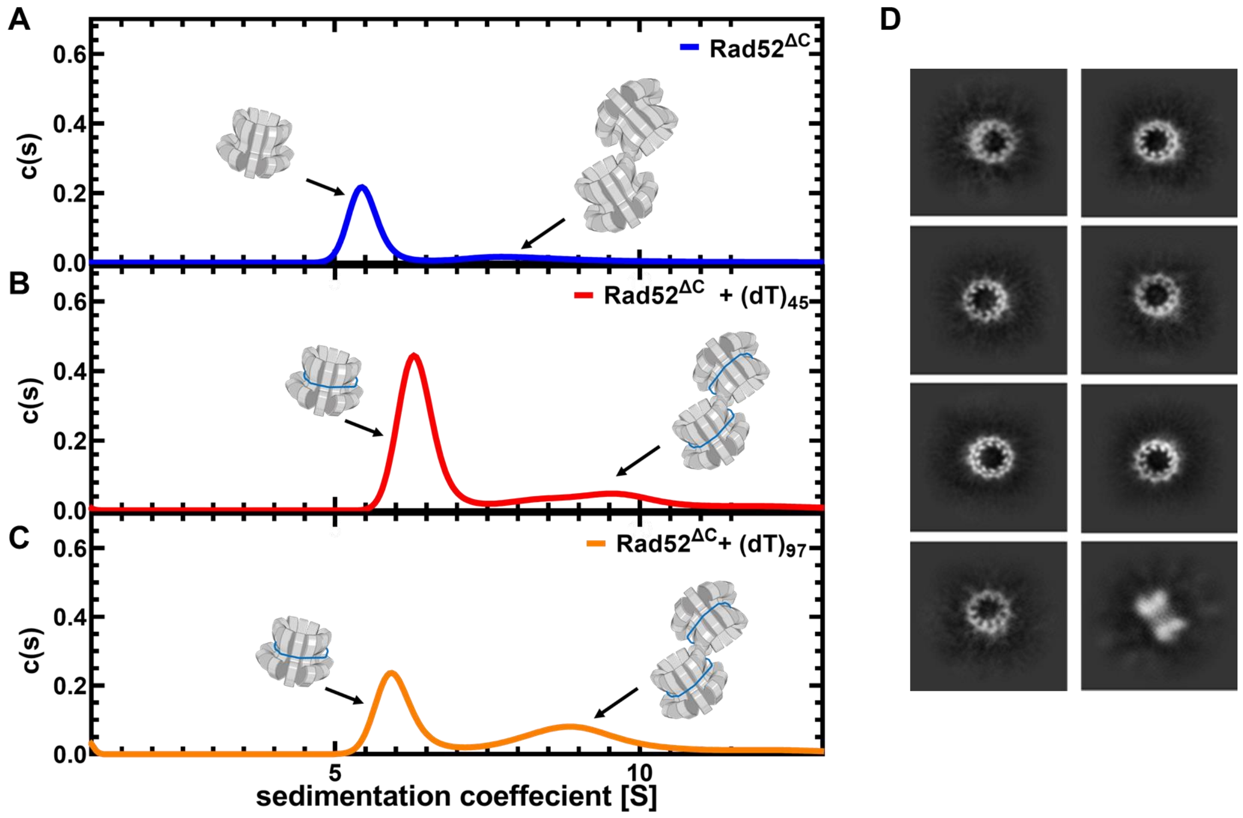

**Supplementary Figure 14. Rad52$^{DC}$ is also a homodecamer in solution.** Analytical ultracentrifugation sedimentation velocity analysis of **A)** Rad52$^{DC}$ in the absence or presence of **B)** (dT)$_{45}$ or **C)** (dT)$_{97}$ show that the N-terminal half of the protein primarily behaves as a single higher order species corresponding to a homodecamer. A small fraction of the double-ring species is also observed. Both species bind to short and longer ssDNA, however, the species distribution does not change. **D)** 2D classes of Rad52$^{DC}$ in cryo-EM analysis show homodecameric rings.

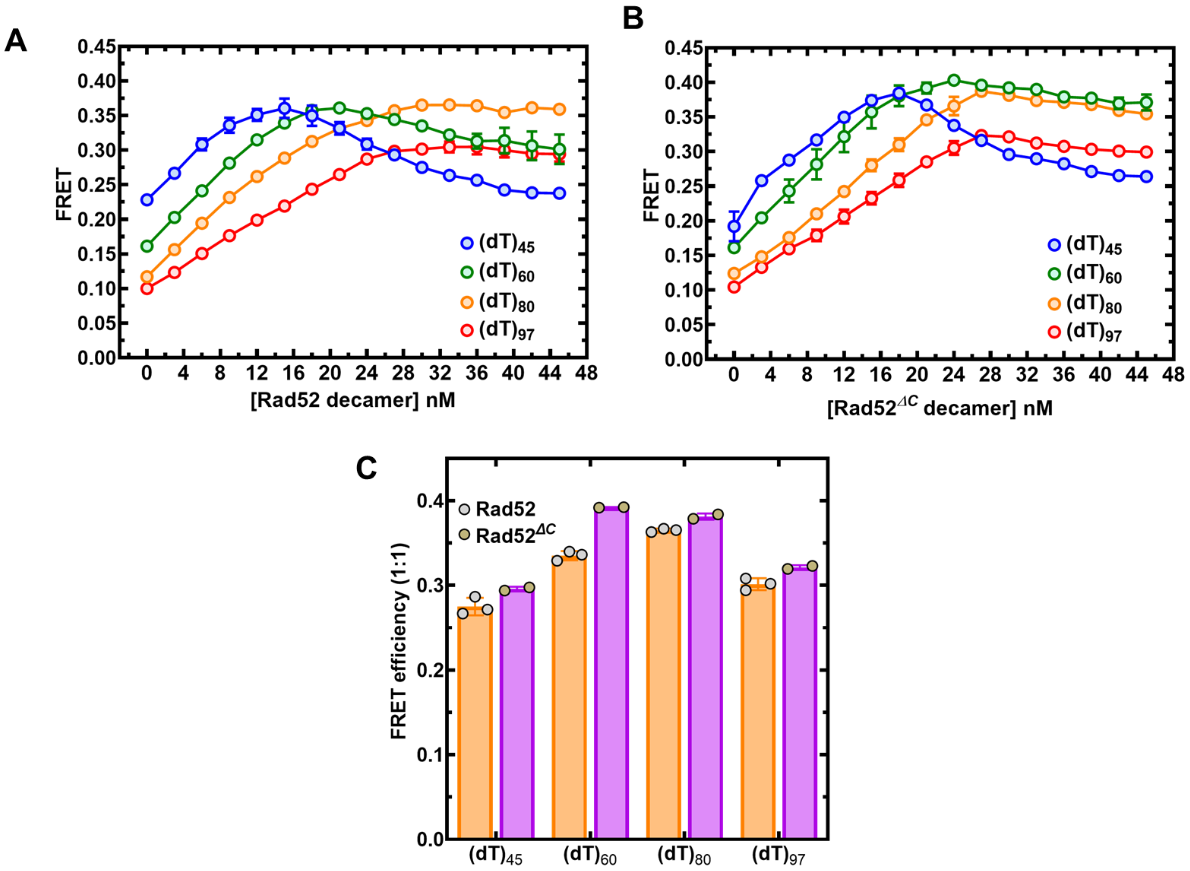

**Supplementary Figure 15. ssDNA length dependent change in Rad52 binding induced FRET.** ssDNA of varying lengths carrying 5′-Cy5 and 3′-Cy3 fluorophores were subject to **A)** Rad52 or **B)** Rad52*DC* binding. Changes in FRET were captured by exciting Cy3 and monitoring changes in Cy5 emission. **C)** Optimal FRET for Rad52 is observed on the $(dT)_{80}$ substrate. For $Rad52^{DC}$, highest FRET efficiency is observed on the $(dT)_{60}$ substrate. These data also reveal that the C-tail influences wrapping of the ssDNA around Rad52.

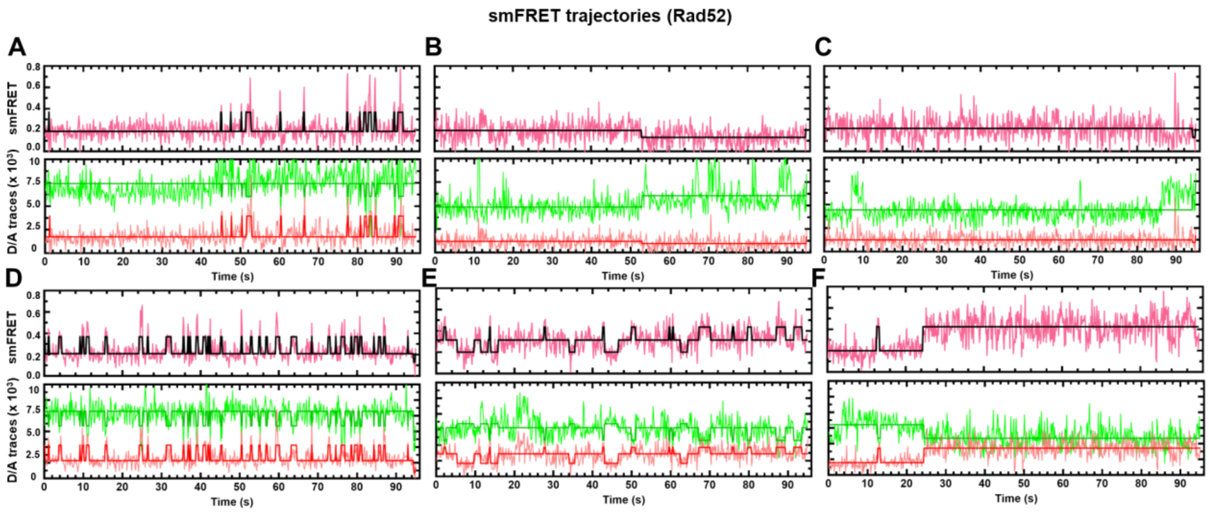

**Supplementary Figure 16. Additional single-molecule FRET trajectories for Rad52.** Trajectories from six different single DNA molecules are shown from the analysis of full length Rad52.

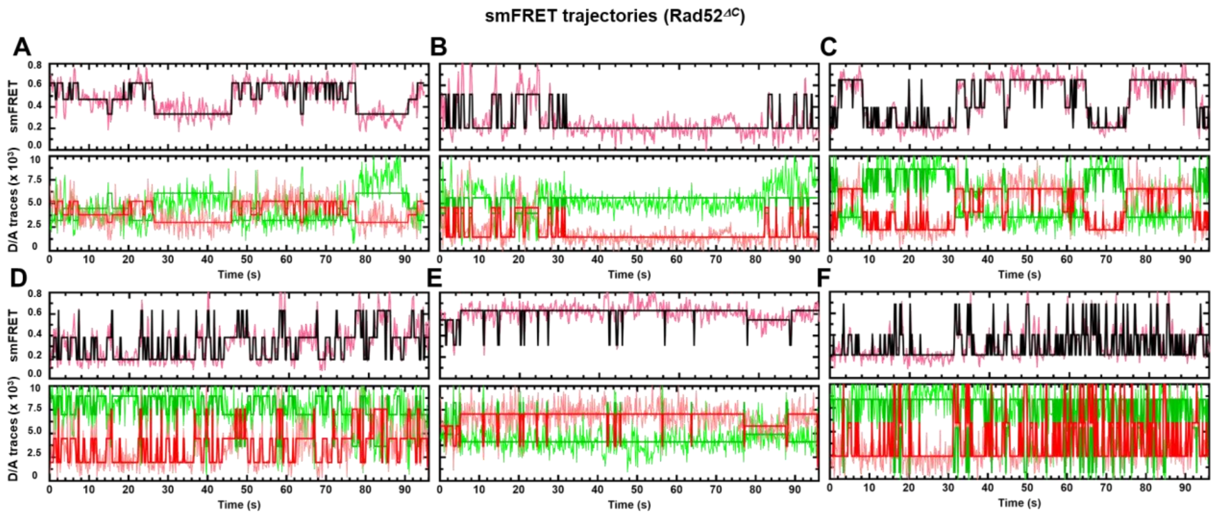

**Supplementary Figure 17. Additional single-molecule FRET trajectories for Rad52$^{DC}$.** Trajectories from six different single DNA molecules are shown from the analysis of Rad52$^{DC}$.

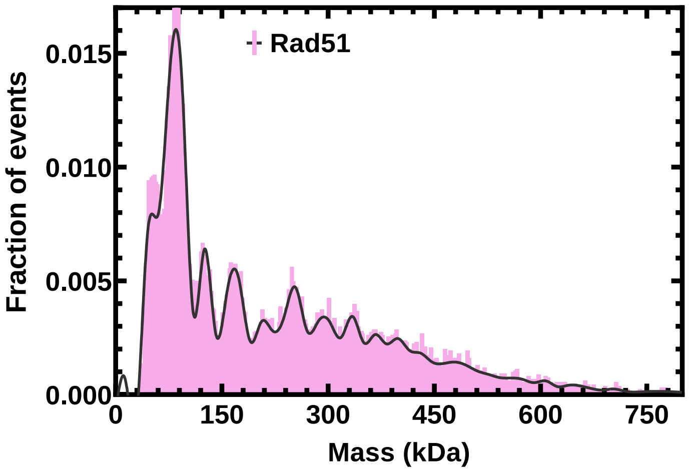

**Supplementary Figure 18. Mass photometry analysis of yeast Rad51.** Mass photometry analysis of *S. cerevisiae* Rad51 shows the presence of multiple oligomeric species.

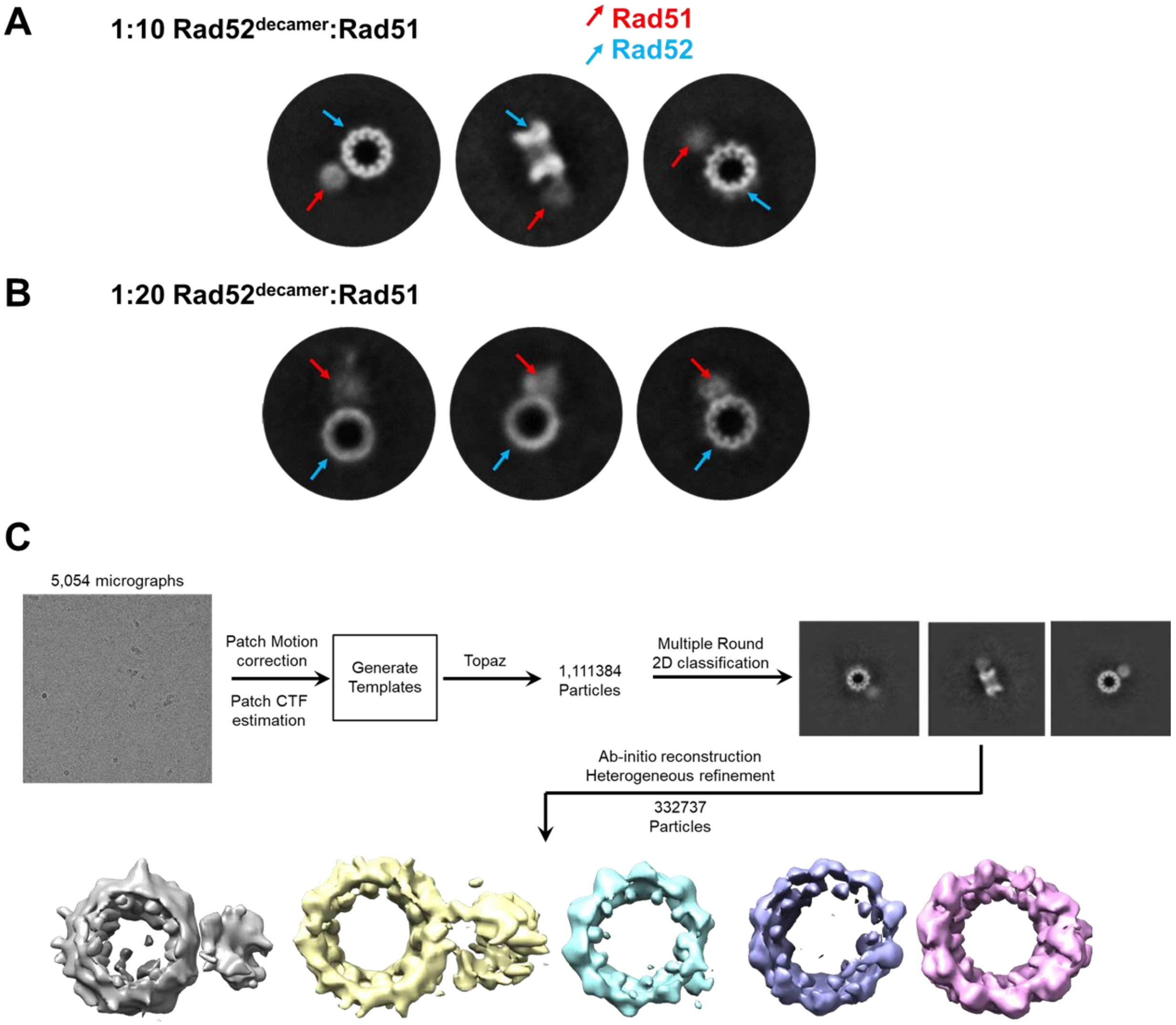

**Supplementary Figure 19. Interaction of Rad52-Rad51 complex.** 2D class averages from the cryo-EM analysis of the Rad51-Rad52 complex when formed using **A)** 1:10 and **B)** 1:20 molar ratios of Rad52 (decamer) to Rad51 are shown. Rad51 molecules are bound to the Rad52 ring, but only at a single defined position. **C)** Workflow of Single particle analysis of Rad52-Rad51 Cryo-EM data

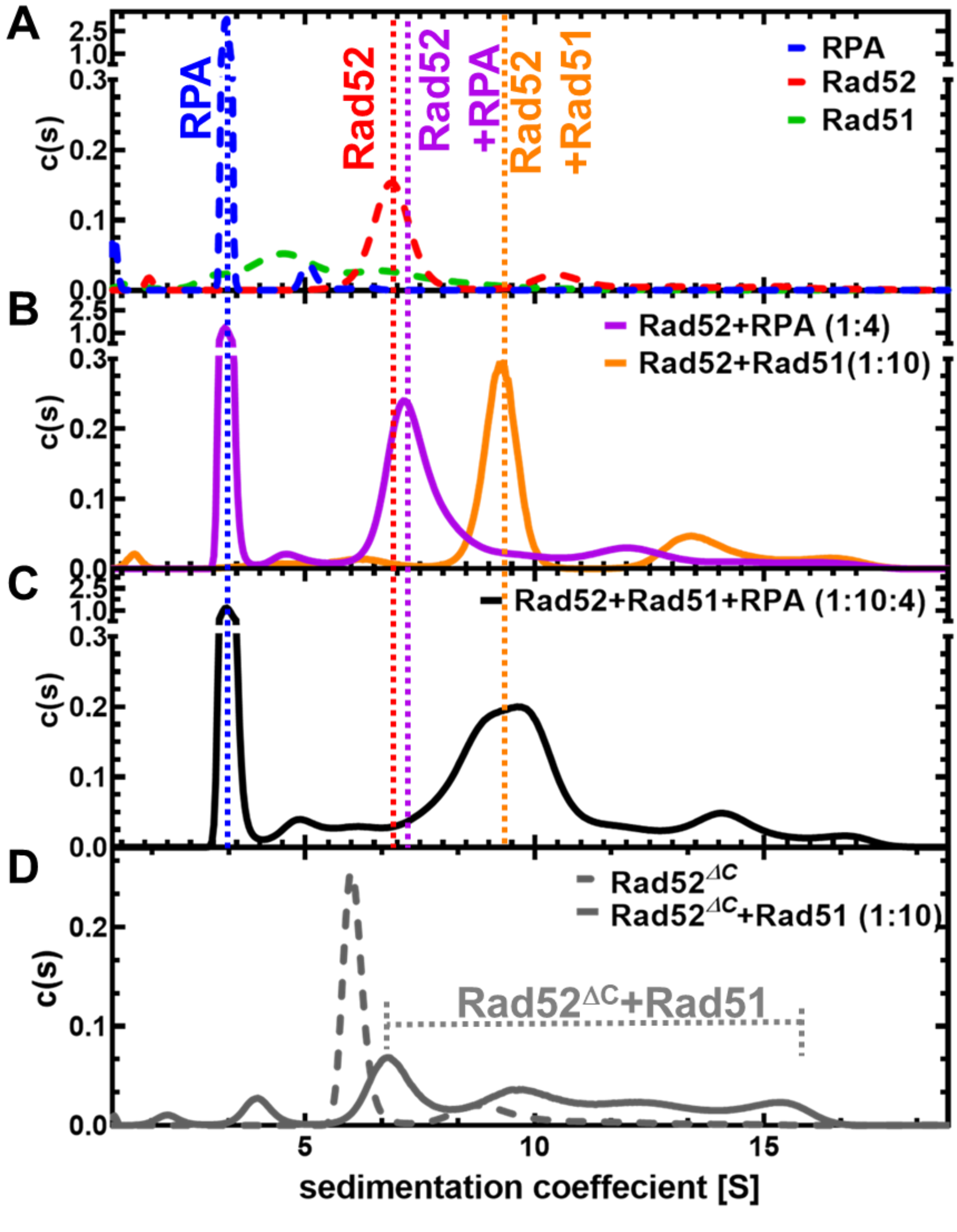

**Supplementary Figure 20. RPA, Rad52, and Rad51 form a stable complex in the absence of ssDNA.** Analytical ultracentrifugation sedimentation velocity analysis of RPA, Rad51, and Rad52 alone and in complex. **A)** Sedimentation profiles of RPA (4 µM), Rad52 (10 µM monomer), and Rad51 (10 µM). RPA is a single heterotrimer. Rad52 is predominantly a homodimer. Rad51 migrates as several polydisperse complexes. **B)** A complex between Rad52 (10 µM) and RPA (4 µM) or Rad51 (10 µM) shows formation of complexes. Even though 10 RPA binding sites are present in the Rad52 decamer, only a few RPA molecules are bound to the Rad52 ring. **C)** When all three proteins are mixed, a large species corresponding to RPA-Rad52-Rad51 are observed. **D)** Rad52$^{DC}$ displays reduced complex formation with Rad51. The sedimentation profiles of Rad52$^{DC}$ or Rad52$^{DC}$+Rad51 (10µM monomer of Rad52$^{DC}$ with or without 10µM Rad51) show a defined Rad52 $^{DC}$ species. In contrast, the Rad52$^{DC}$+Rad51 complex displays more polydisperse behavior suggesting non-uniform complex formation.

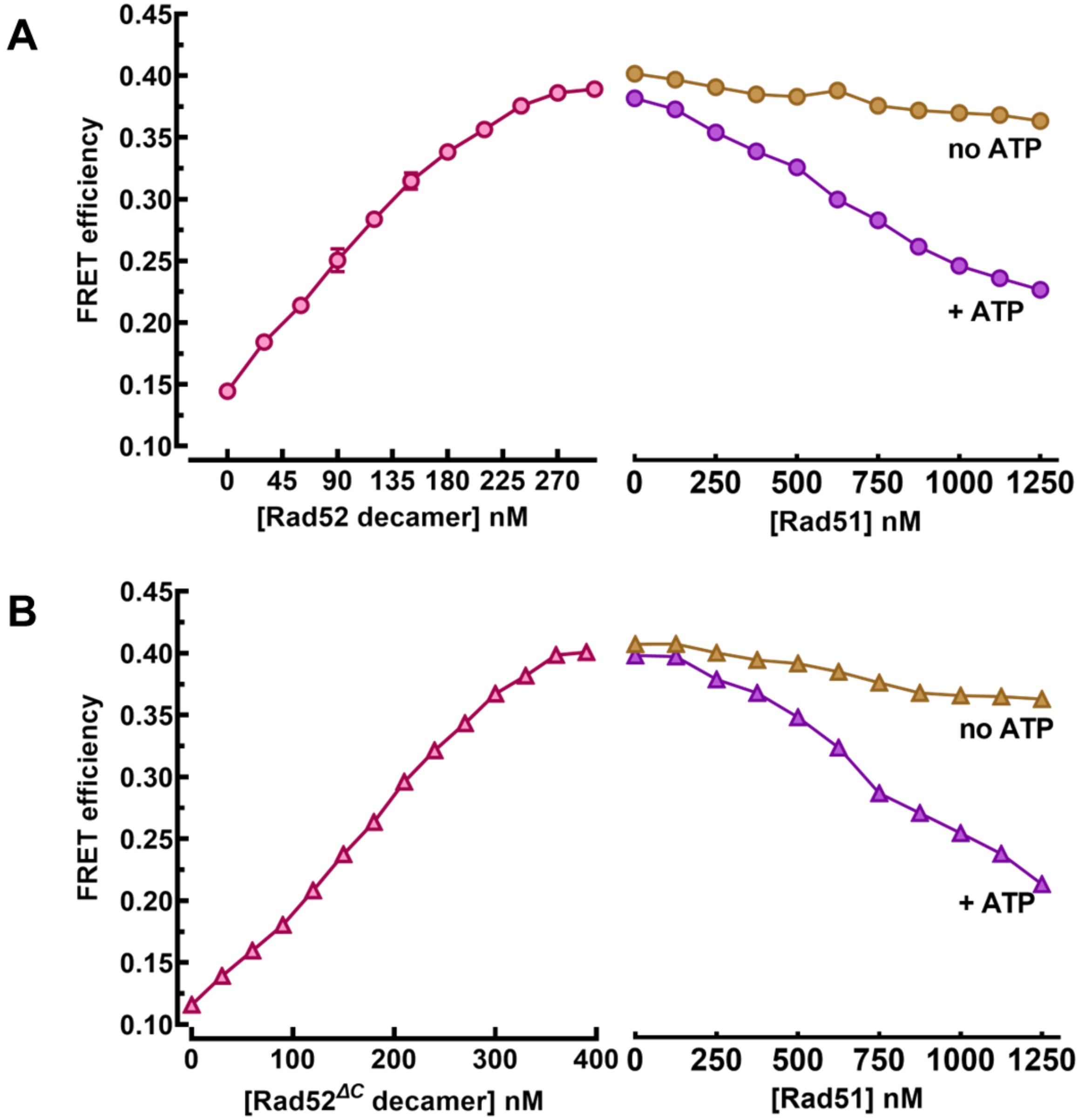

**Supplementary Figure 21. Ensemble FRET measurements of Rad51-binding to Cy3-Cy5 end labeled (dT)$_{80}$ ssDNA. A)** Rad52 or **B)** Rad52$^{DC}$ were prebound to a Cy3-(dT)$_{80}$-Cy5 substrate and Rad51 was introduced in the absence or presence of ATP. Data shows that both Rad52 and Rad52$^{DC}$-DNA complexes can be remodeled by Rad51 when sufficient time is provided to reach equilibrium.

**Supplementary Table 1: Mass Photometer mass distribution.**

| $(dT)_{84}$ kDa | Rad52 (n=8) kDa | Rad52 + $(dT)_{84}$ (n = 11) kDa | $Rad52^{\Delta C}$ (n=3) kDa | $Rad52^{\Delta C}$ + $(dT)_{84}$ (n=3) kDa |
|---|---|---|---|---|
| 63 ±19 | 72.9±14 | 87±26. | 66±33.5 | 97 |
| - | 498.4±24 | 522.5±25 | 255±12 | 293±10 |
| - | 991.8±75 | 1078±68 | 517±29 | 606±17 |
| - | 1551±99 | 1562±37 | 782±46 | 931.5±29 |

** 1) Major peak (>85% of the population) is highlighted in bold*

*2) All the different Rad52 and $Rad52^{DC}$ peaks show ssDNA binding as can be seen in the systematic shift in the entire distribution*

*3) The $(dT)_{84}$ mass estimation is prone to lot of error because the actual mass of the ssDNA is 25,490.5 gm/mole which is at/below the detection limit of the of the scope*

**Supplementary Table 2. CryoEM data collection and model statistics**

| | Rad52 Apo |
|---|---|
| EMD | 29695 |
| PDB | 8G3G |
| **Data Collection and processing** | |
| Magnification | 150,000x |
| Voltage (kV) | 200 |
| Electron exposure ($e^-$/ Å$^2$) | 50.7 |
| Defocus range (mm) | -0.8 to -2.4 |
| Nominal Pixel Size (Å) | 0.94 |
| Symmetry imposed | C10 |
| Initial Particle images | 701,962 |
| Final Particle images | 180,545 |
| Map resolution (Å) | 3.3/3.4/3.8 |
| FSC threshold | 0/0.143/0.5 |
| Map resolution range (Å) | 2.8 to 6.0 |
| **Refinement** | |
| Model resolution (Å) | 3.5 |
| FSC threshold | 0.143 |
| Map sharpening B-factor (Å2) | |
| Model composition | |
| Chains | 10 |
| Non-hydrogen atoms | 6930 |
| Protein residues | 890 |
| Ligands | 0 |
| Water | 0 |
| Average B-factors (Å$^2$) | |
| Protein | 71.88 |
| Bond RMSD | |
| Bond lengths (Å) | 0.003 |
| Bond angles (°) | 0.719 |
| **Validation** | |
| MolProbity score | 1.49 |
| All-atom clash score | 8.27 |
| Ramachandran plot | |
| Favored (%) | 97.87 |
| Allowed (%) | 2.18 |
| Outliers (%) | 0 |

**Supplementary Table 3: DNA oligonucleotides used in various assays.**

| Name | Sequence (5′ to 3′) | Comments |
|---|---|---|
| 5′-FAM-(dT)$_{35}$ | 5'- /56-FAM/TTT TTT TTT TTT TTT TTT TTT TTT TTT TTT TTT TT -3' | Fluorescence Anisotropy |
| 5′-FAM-(dT)$_{70}$ | 5'- /56-FAM/TTT TTT TTT TTT TTT TTT TTT TTT TTT TTT TTT TTT TTT TTT TTT TTT TTT TTT TTT TTT TTT TTT TTT T -3' | Fluorescence Anisotropy |
| 5′-FAM-32bp-duplex | 5'-/56-FAM/CGA ATG TAA CCA TCG TTG GTC GGC AGC AGG GC -3' | 5′-FAM labelled oligo (sequence shown on the left) was hybridized with an unlabeled complementary strand to form a 32 base paired duplex for Rad52 binding experiments |
| (dT)$_{20}$ | 5'- TTT TTT TTT TTT TTT TTT TT -3' | Trp quenching |
| (dT)$_{30}$ | 5'- TTT TTT TTT TTT TTT TTT TTT TTT TTT TTT -3' | Trp quenching |
| (dT)$_{45}$ | 5'- TTT TTT TTT TTT TTT TTT TTT TTT TTT TTT TTT TTT TTT TTT TTT -3' | Trp quenching |
| (dT)$_{60}$ | 5'- TTT TTT TTT TTT TTT TTT TTT TTT TTT TTT TTT TTT TTT TTT TTT TTT TTT TTT TTT TTT -3' | Trp quenching |
| (dT)$_{84}$ | 5'- TTT TTT TTT TTT TTT TTT TTT TTT TTT TTT TTT TTT TTT TTT TTT TTT TTT TTT TTT TTT TTT TTT TTT TTT TTT TTT TTT TTT -3' | Trp quenching, Mass Photometry |
| (dT)$_{97}$ | 5'- TTT TTT TTT TTT TTT TTT TTT TTT TTT TTT TTT TTT TTT TTT TTT TTT TTT TTT TTT TTT TTT TTT TTT TTT TTT TTT TTT TTT TTT TTT TTT TTT T -3' | Cryo-EM substrate, AUC |
| poly(dT) | - | Mean oligo length is 1100 nt |
| 5′-Cy5-(dT)$_{45}$-Cy3-3′ | 5'- /5Cy5/TTT TTT TTT TTT TTT TTT TTT TTT TTT TTT TTT TTT TTT TTT TTT /3Cy3Sp/ -3' | Fluorometer based ssDNA wrapping FRET assay |
| 5′-Cy5-(dT)$_{60}$-Cy3-3′ | 5'- /5Cy5/TTT TTT TTT TTT TTT TTT TTT TTT TTT TTT TTT TTT TTT TTT TTT TTT TTT TTT TTT TTT /3Cy3Sp/ -3' | Fluorometer based ssDNA wrapping FRET assay |
| 5′-Cy5-(dT)$_{80}$-Cy3-3′ | 5'- /5Cy5/TTT TTT TTT TTT TTT TTT TTT TTT TTT TTT TTT TTT TTT TTT TTT TTT TTT TTT TTT TTT TTT TTT TTT TTT TTT TT/3Cy3Sp/ -3' | Fluorometer based ssDNA wrapping FRET assay |
| 5′-Cy5-(dT)$_{97}$-Cy3-3′ | 5'- /5Cy5/TTT TTT TTT TTT TTT TTT TTT TTT TTT TTT TTT TTT TTT TTT TTT TTT TTT TTT TTT TTT TTT TTT TTT TTT TTT TTT TTT TTT TTT TTT TTT TTT T/3Cy3Sp/ -3' | Fluorometer based ssDNA wrapping FRET assay |

| | | |
|---|---|---|
| 5' -Cy5-18mer-Bio-3' | 5' /5Cy5/GCC TCG CTG CCG TCG CCA/3Bio/ 3' | Biotin-ssDNA handle with 5'- Cy5 |
| 5' - 18mer-$(dT)_{40}$-Cy3-3' | 5'- TGG CGA CGG CAG CGA GGC $(dT)_{40}$ /3Cy3Sp/ -3' | Partial duplex with $(dT)_{40}$ overhang |
| 5' - 18mer-$(dT)_{60}$-Cy3-3' | 5'- TGG CGA CGG CAG CGA GGC $(dT)_{60}$ /3Cy3Sp/ -3' | Partial duplex with $(dT)_{60}$ overhang |
| 5' - 18mer-$(dT)_{80}$-Cy3-3' | 5'- TGG CGA CGG CAG CGA GGC $(dT)_{80}$ /3Cy3Sp/ -3' | Partial duplex with $(dT)_{80}$ overhang |

### Supplementary Table 4: DNA Primers used for Rad52 mutagenesis.

| Name | Sequence (5′ to 3′) | Comments |
|---|---|---|
| Rad52$^{1\text{-}212}$ F | 5'-ATATACATATGAATGAAATTATGGATATGGATGAG-3' | Rad52$^{DC}$ |
| Rad52$^{1\text{-}212}$ R | 5'-GCAGTACTCGAGCTTCGTCGAGTCGGGATTG-3' | Rad52$^{DC}$ |
| Rad52$^{213\text{-}471}$ F | 5'-ATATACATATGAACCTGGTGAAAATAGAAAATACAGTAAGTCG-3' | Rad52$^{DN}$ |
| Rad52$^{213\text{-}471}$ R | 5'-GCAGTACTCGAGTCAAGTAGGCTTGCGTGCA-3' | Rad52$^{DN}$ |
| Rad52$^{293\text{-}471}$ F | 5'-ATATACATATGGATCCCGTTGTAGCTAAGCAAAGCCC-3' | Rad52$^{DN*}$ |
| Rad52$^{293\text{-}471}$ R | 5'-GCAGTACTCGAGTCAAGTAGGCTTGCGTGCATGCA-3' | Rad52$^{DN*}$ |
